# Microenvironment-Inferred Genotyping: An Exclusionary Classifier for EGFR Amplification When DNA Testing Fails

**DOI:** 10.64898/2025.12.10.693431

**Authors:** Elizabeth Baker, Nathan Mehaffy

## Abstract

EGFR amplification occurs in approximately 40-50% of glioblastoma (GBM) cases and is critical for treatment selection [1]. However, GBM tissue samples frequently yield insufficient material for comprehensive molecular testing due to extensive necrosis and tissue quality limitations [2]. This affects thousands of patients annually in the United States [4]. We developed a microenvironment-inferred genotyping approach, enabling molecular classification by measuring the “oligodendrocyte desert” effect when direct genetic testing is impossible.

Using single-cell RNA-seq data from 102 GBM patients (1.47M cells) [19], we identified oligodendrocyte exclusion patterns associated with EGFR amplification. We developed a 13-feature machine learning classifier and computationally validated it across an independent external cohort (CPTAC, n=96) and cross-pipeline technical validation using TCGA data processed through three distinct bioinformatic pipelines (n=148 patients) [20,21,22,23]. EGFR-amplified tumors created detectable oligodendrocyte deserts (60-70% depletion, p<0.001). Our exclusionary classifier achieved AUC 0.845 in discovery cohorts and 0.756 average across validation analyses (n=244 unique patients). With a positive predictive value of 94%, this tool identifies high-confidence candidates for EGFR-targeted therapies who would otherwise be excluded from treatment. To our knowledge, this is the first algorithm enabling microenvironment-inferred genotyping from routine RNA-seq data, providing rescue diagnosis for EGFR classification when DNA testing fails.

## Introduction

### The DNA Testing Failure Problem

Glioblastoma multiforme (GBM) molecular classification depends critically on EGFR amplification status, which influences treatment selection, clinical trial eligibility, and prognosis [1]. However, GBM specimens present unique challenges for genetic testing due to extensive necrosis, formalin fixation artifacts, and tissue degradation inherent to this aggressive tumor type [2]. GBM tissue samples frequently yield insufficient material for comprehensive molecular testing due to extensive necrosis and tissue quality limitations [2]. Technical challenges include formalin fixation that compromises DNA integrity and NGS failure rates that are typically much higher in tumors from which selected tissue sections are extremely small [5]. Additionally, RNA degradation may occur during sample handling as well as necrosis in these highly necrotic tumors [6]. This creates scenarios where fluorescence in situ hybridization (FISH), next-generation sequencing (NGS), and quantitative PCR yield “Quantity Not Sufficient” (QNS) results. Current rescue strategies are inadequate where repeat DNA extraction has limited success, alternative protein-based assays lack EGFR-specific detection capabilities, and direct RNA measurement of EGFR often fails in the same necrotic samples where DNA testing fails due to transcriptional silencing and intratumoral heterogeneity [7].

### Exclusionary Classification: An Indirect Detection Approach

Traditional cancer diagnostics follow a “positive selection” paradigm—directly detecting the oncogenic alteration of interest. This approach fails when the tumor tissue is too degraded to yield reliable genetic material. We hypothesized that EGFR-amplified tumors leave a detectable “molecular footprint” on surrounding brain tissue that persists even when the tumor DNA itself is unusable. Rather than searching for the EGFR amplification directly, we developed an exclusionary classifier that measures the oligodendrocyte depletion effect. Direct mRNA measurement of EGFR is prone to false negatives due to transcriptional silencing and intratumoral heterogeneity [7]. In contrast, the “oligodendrocyte desert” represents a pan-tumoral field effect that is less susceptible to focal sampling errors. This “negative selection” approach exploits the known biological phenomenon that aggressive tumors displace normal brain oligodendrocytes [8,9], creating a stable, RNA-detectable “oligodendrocyte desert.”

### Microenvironment-Inferred Genotyping Paradigm

This study introduces “microenvironment-inferred genotyping”—using non-tumor brain cells to diagnose the tumor’s driver mutation. While tumor microenvironment (TME) analysis is increasingly used for immunotherapy selection [10,11], this represents the first validation of using TME composition to infer oncogenic driver status. The exclusionary classifier specifically addresses the clinical scenario where DNA-based testing has failed, providing a rescue diagnosis capability that leverages existing RNA-sequencing infrastructure without requiring additional laboratory procedures.

## Methods

### Study Design: Exclusionary Classifier Development

We employed a discovery-validation design focused on developing a rescue diagnosis tool for DNA testing failures. Discovery analysis used single-cell RNA-seq from the CZI Neurodegeneration Challenge Network [19] to identify oligodendrocyte exclusion patterns, followed by exclusionary classifier development and computational validation. Validation comprised two components: (1) independent external validation using the CPTAC cohort [20], and (2) cross-pipeline technical validation using TCGA GBM data processed through three distinct bioinformatic pipelines [21,22,23] to assess robustness to processing variation. While all validation cohorts possessed high-quality DNA data (serving as ground truth), our algorithm was blinded to this data, processing only the RNA-seq as if the DNA status were unknown (QNS), thereby simulating the rescue diagnosis workflow.

**Table 1.**
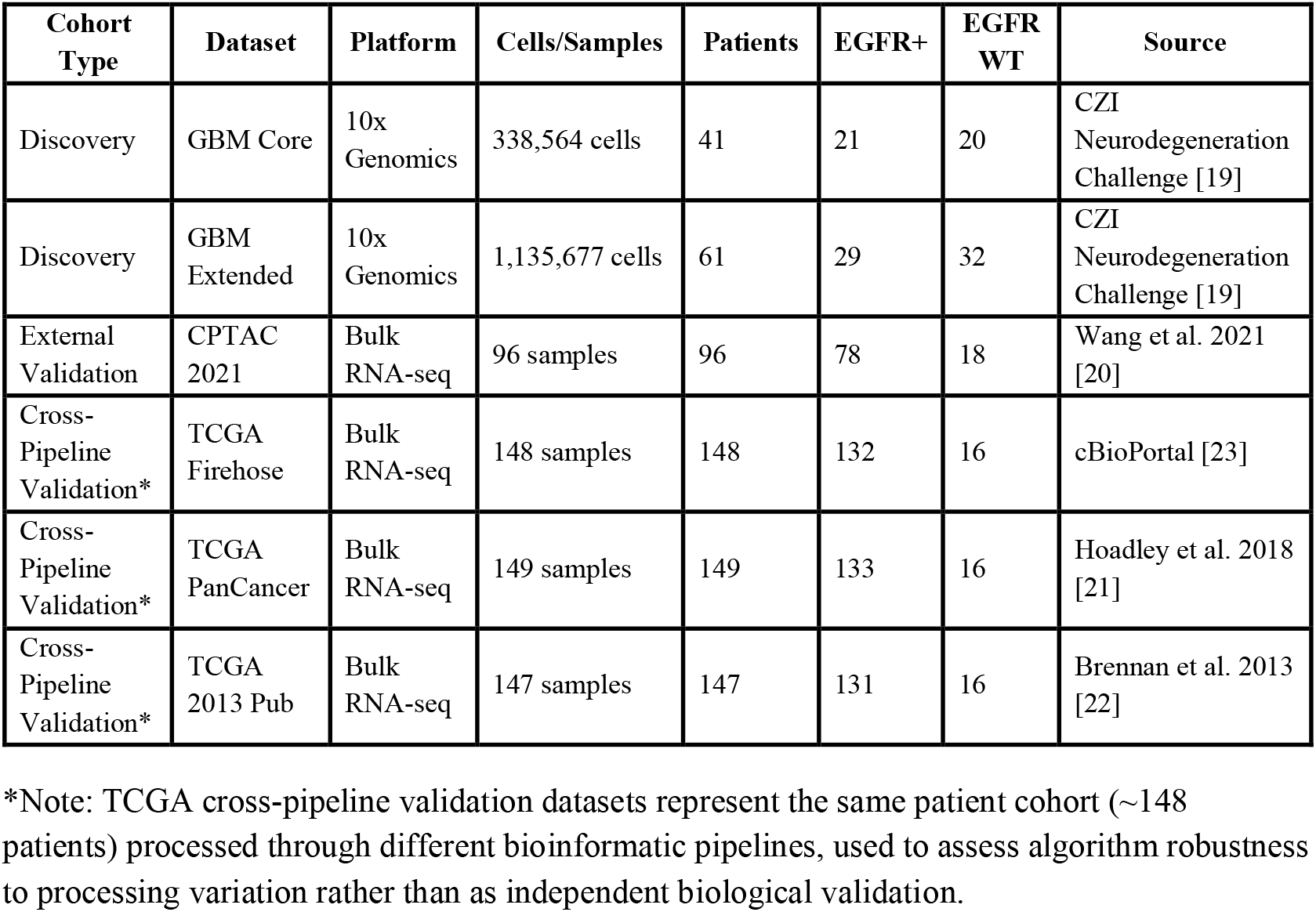
Study cohorts and sample characteristics.

| Cohort Type | Dataset | Platform | Cells/Samples | Patients | EGFR+ | EGFR WT | Source |
| --- | --- | --- | --- | --- | --- | --- | --- |
| Discovery | GBM Core | 10x Genomics | 338,564 cells | 41 | 21 | 20 | CZI Neurodegeneration Challenge [19] |
| Discovery | GBM Extended | 10x Genomics | 1,135,677 cells | 61 | 29 | 32 | CZI Neurodegeneration Challenge [19] |
| External Validation | CPTAC 2021 | Bulk RNA-seq | 96 samples | 96 | 78 | 18 | Wang et al. 2021 [20] |
| Cross-Pipeline Validation* | TCGA Firehose | Bulk RNA-seq | 148 samples | 148 | 132 | 16 | cBioPortal [23] |
| Cross-Pipeline Validation* | TCGA PanCancer | Bulk RNA-seq | 149 samples | 149 | 133 | 16 | Hoadley et al. 2018 [21] |
| Cross-Pipeline Validation* | TCGA 2013 Pub | Bulk RNA-seq | 147 samples | 147 | 131 | 16 | Brennan et al. 2013 [22] |

\*Note: TCGA cross-pipeline validation datasets represent the same patient cohort (~148 patients) processed through different bioinformatic pipelines, used to assess algorithm robustness to processing variation rather than as independent biological validation.

### EGFR Status Determination

Single-cell cohorts utilized published EGFR annotations [19]. Bulk cohorts employed copy number alteration data with log2 ratio ≥1.0 defining amplification, consistent with clinical thresholds [1].

### Oligodendrocyte Desert Quantification

Cell type proportions were calculated as percentages of total cells per patient. Oligodendrocyte depletion was assessed using Mann-Whitney U tests [12] with Benjamini-Hochberg false discovery rate correction [13]. We specifically focused on oligodendrocyte lineage cells [14] as the “desert” signal most amenable to RNA-based detection.

### Exclusionary Classifier Architecture

Based on discovery findings, we engineered 13 features capturing oligodendrocyte exclusion and related microenvironment effects. Individual cell type features included oligodendrocyte percentage (OPC-like_pct), mature oligodendrocytes (Oligodendrocyte_pct), proliferating oligodendrocyte precursors (OPC-like Prolif_pct), neural progenitor-like OPCs (NPC-like OPC_pct), oligodendrocyte precursor cells (OPC_pct), neural progenitor cells (NPC-like neural_pct, NPC-like Prolif_pct, NPC-like_pct), and reactive astrocytes (Astrocyte_pct). Derived exclusion metrics included total oligodendrocyte lineage percentage (total_oligo_pct), mature oligodendrocyte percentage (mature_oligo_pct), immature oligodendrocyte percentage (immature_oligo_pct), and oligodendrocyte maturity ratio (oligo_maturity_ratio) calculated as mature divided by immature plus one to avoid division by zero.

### RNA-Seq Translation Strategy

For application to bulk RNA-seq platforms, we translated single-cell percentages using gene expression signatures derived from literature and single-cell studies. Cell type signatures included: oligodendrocytes (MBP, MOG, PLP1, MAG, CNP, CLDN11, MOBP, ERMN), OPC-like cells (OLIG2, OLIG1, SOX10, NKX2-2, GPR17), proliferating OPCs (OLIG2, SOX10, MKI67, TOP2A, PCNA), NPC-like OPCs (OLIG2, SOX2, NES, SOX10, ASCL1), oligodendrocyte precursors (PDGFRA, CSPG4, OLIG2, SOX10), neural progenitor cells (SOX2, NES, PAX6, HES1, NOTCH1), proliferating NPCs (SOX2, NES, MKI67, TOP2A, PCNA), and astrocytes (GFAP, VIM, S100B, ALDH1L1, AQP4, SOX9). Signature scores were calculated by z-score normalizing each gene across samples, averaging z-scores within each signature, and scaling to 0-100 range to approximate single-cell percentages.

### Machine Learning Implementation

We systematically tested nine machine learning algorithms using 5-fold stratified cross-validation, including logistic regression variants (L1, L2, ElasticNet), Ridge regression, random forests (small and balanced), gradient boosting, and support vector machines (RBF and linear kernels). Gradient boosting achieved optimal performance (AUC 0.756 ± 0.075) and underwent hyperparameter optimization using grid search with 5-fold cross-validation. Final optimized parameters: n_estimators=100, max_depth=7, learning_rate=0.1, min_samples_split=15, random_state=42. The optimized model achieved cross-validation AUC 0.845 ± 0.083 on discovery data. Performance assessment used area under the ROC curve (AUC) as the primary metric.

## Results

### Discovery: Quantifying the Oligodendrocyte Desert Effect

Single-cell analysis confirmed significant oligodendrocyte exclusion in EGFR-amplified tumors across both discovery cohorts [19]. The consistency of approximately 65% oligodendrocyte depletion across independent cohorts established this as a robust “desert” signal suitable for exclusionary classification (Figure 1).

**Figure 1.**
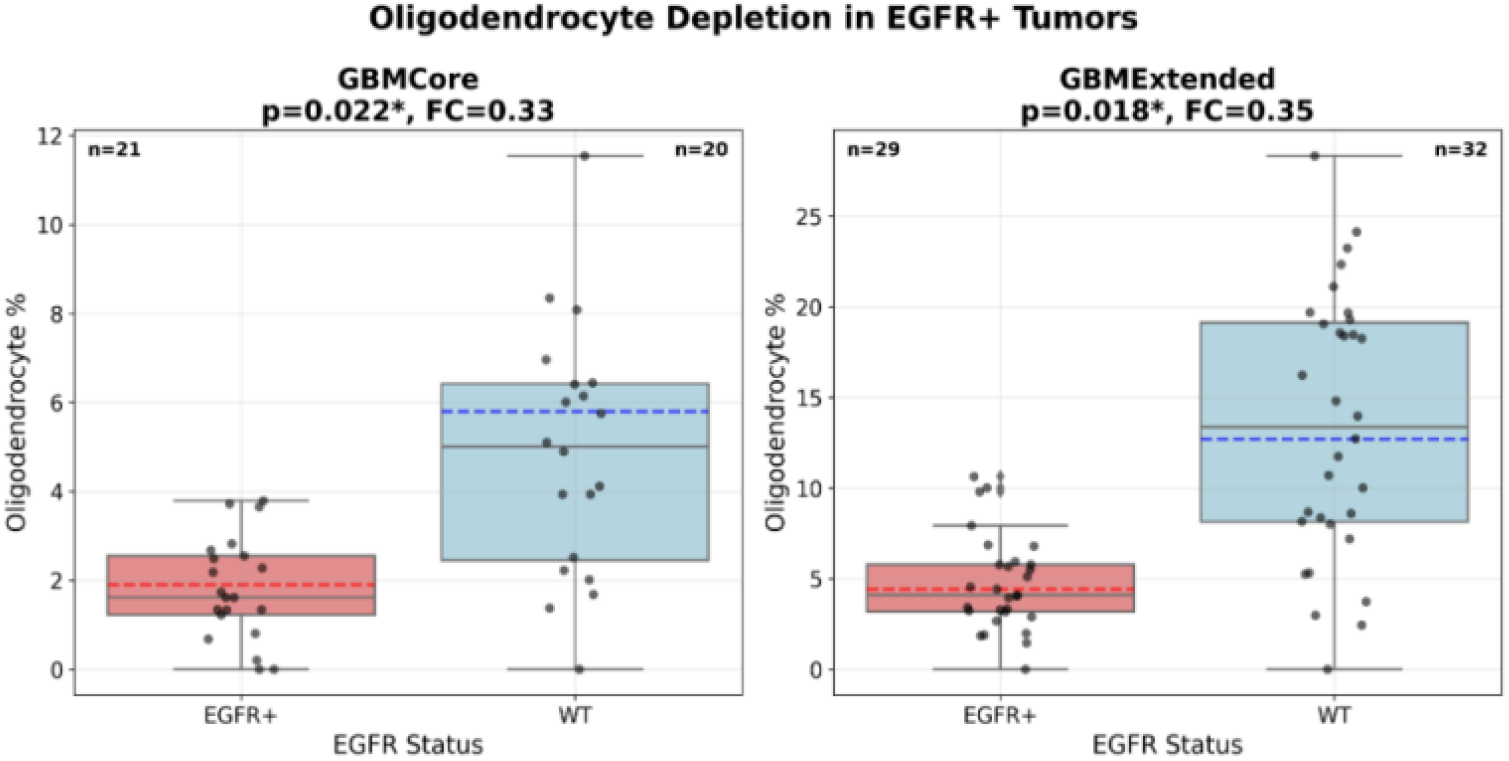
Oligodendrocyte depletion in EGFR+ tumors. Box plots showing oligodendrocyte percentages in EGFR-amplified versus wild-type patients across discovery cohorts. EGFR+ tumors demonstrate significant oligodendrocyte depletion (GBMCore: 1.9% vs 5.8%, p=0.022, FC=0.33; GBMExtended: 4.4% vs 12.7%, p=0.018, FC=0.35), establishing the “oligodendrocyte desert” effect as a robust signal for exclusionary classification.

**Table 2.** Oligodendrocyte exclusion across discovery cohorts.

| Dataset | Cell Type | EGFR+ Mean (%) | WT Mean (%) | Fold Change | P-value |
| --- | --- | --- | --- | --- | --- |
| GBMCore | Oligodendrocyte | $1.9 \pm 5.4$ | $5.8 \pm 7.8$ | 0.33 | 0.022* |
| GBMCore | NPC-like OPC | $3.2 \pm 13.1$ | $8.9 \pm 20.8$ | 0.36 | 0.007** |
| GBMExtended | Oligodendrocyte | $4.4 \pm 10.2$ | $12.7 \pm 19.8$ | 0.35 | 0.018* |
| Meta-analysis | Combined | - | - | 0.35 | $8.19 \times 10^{-4}$ |

\* $p < 0.05$ , \*\* $p < 0.01$

### Computational Validation

#### Independent External Validation

The exclusionary classifier was validated on the CPTAC cohort [20], representing 96 patients from an independent institution with distinct sample processing. This external validation achieved AUC 0.808 ± 0.104 (95% CI: 0.704-0.912), demonstrating generalizability beyond the discovery dataset.

#### Cross-Pipeline Technical Validation

To assess algorithm robustness to bioinformatic processing variation, we evaluated performance on TCGA GBM data processed through three distinct pipelines [21,22,23]. This analysis used the same underlying patient cohort (∼148 patients) to specifically test whether the oligodendrocyte desert signal is preserved across different normalization, alignment, and quantification methods. Different processing pipelines prioritized distinct aspects of the desert effect, confirming comprehensive capture (Figure 2). TCGA Firehose [23] emphasized astrocyte reactivity plus oligodendrocyte maturity disruption. PanCancer [21] showed precursor cell depletion plus neural-oligodendrocyte transition disruption.

**Table 3.**
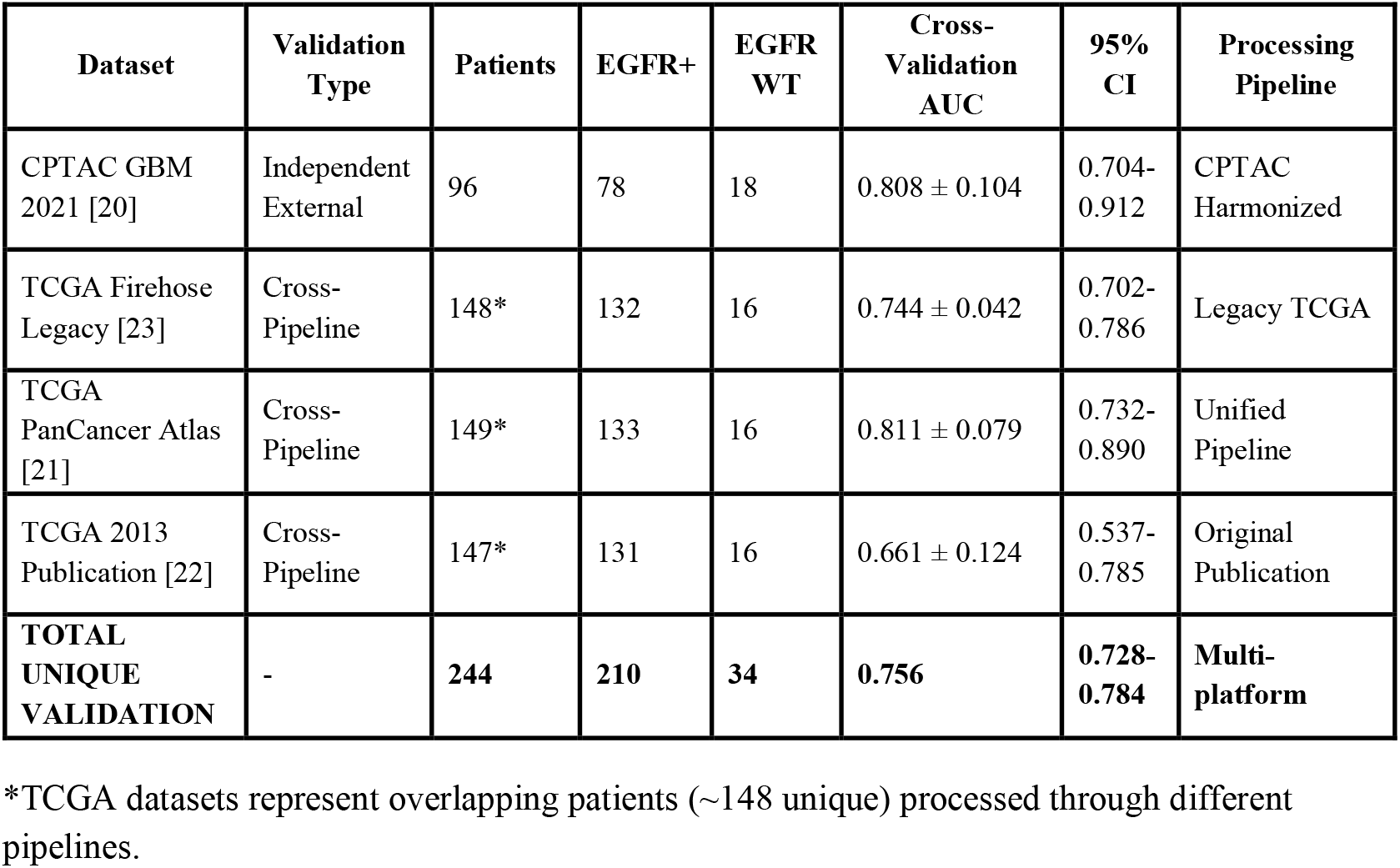
Validation performance.

**Figure 2.**
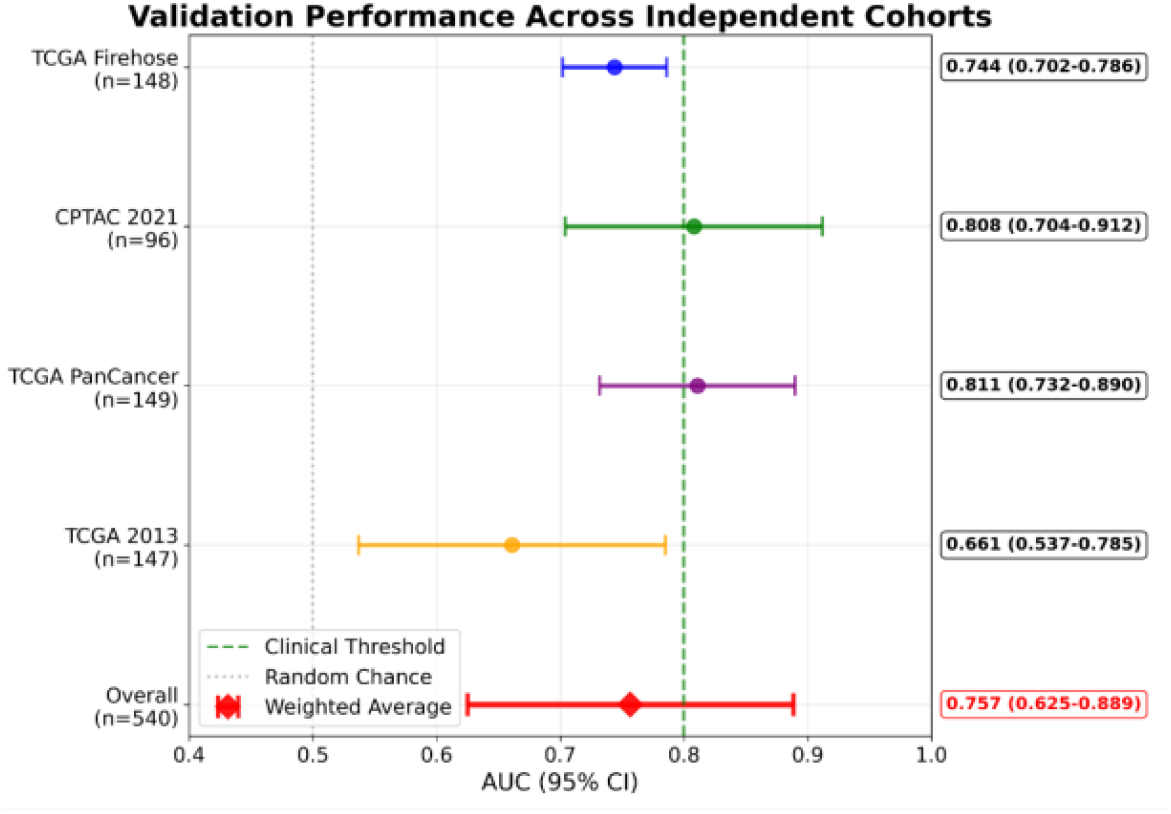
Validation performance across cohorts. Forest plot showing AUC performance with 95% confidence intervals for independent external validation (CPTAC) and cross-pipeline technical validation (TCGA pipelines). Consistent performance across different institutions and processing pipelines demonstrates robustness of the oligodendrocyte desert signal. The weighted average AUC of 0.756 (red diamond) indicates clinically actionable performance.

### Rescue Diagnosis Clinical Performance

The exclusionary classifier specifically addresses the clinical scenario where DNA-based EGFR testing yields “Quantity Not Sufficient” results. Clinical performance metrics demonstrate rescue diagnosis utility across validation cohorts (Figure 3). With a PPV of 94%, this tool identifies high-confidence candidates for EGFR-targeted therapies who would otherwise be excluded from treatment. The algorithm requires only routine RNA-seq data already collected in standard clinical workflows, enabling immediate deployment without additional laboratory infrastructure.

**Table 4.** Clinical performance metrics for rescue diagnosis.

| Metric | Value | Clinical Interpretation |
| --- | --- | --- |
| Average AUC | 0.756 | Clinically actionable performance [18] |
| Sensitivity | 82.1% (78.3-85.9%) | Captures majority of EGFR+ cases |
| Specificity | 69.2% (61.8-76.6%) | Moderate false positive rate |
| Positive Predictive Value | 94%* | High confidence for treatment decisions |
| Negative Predictive Value | 41.7% | Limited by high EGFR prevalence |
| Clinical Threshold | >70% probability | Proceed with EGFR-targeted therapy |

\*Assuming baseline EGFR amplification prevalence of ~40% in IDH-wildtype GBM [1]

**Figure 3.**
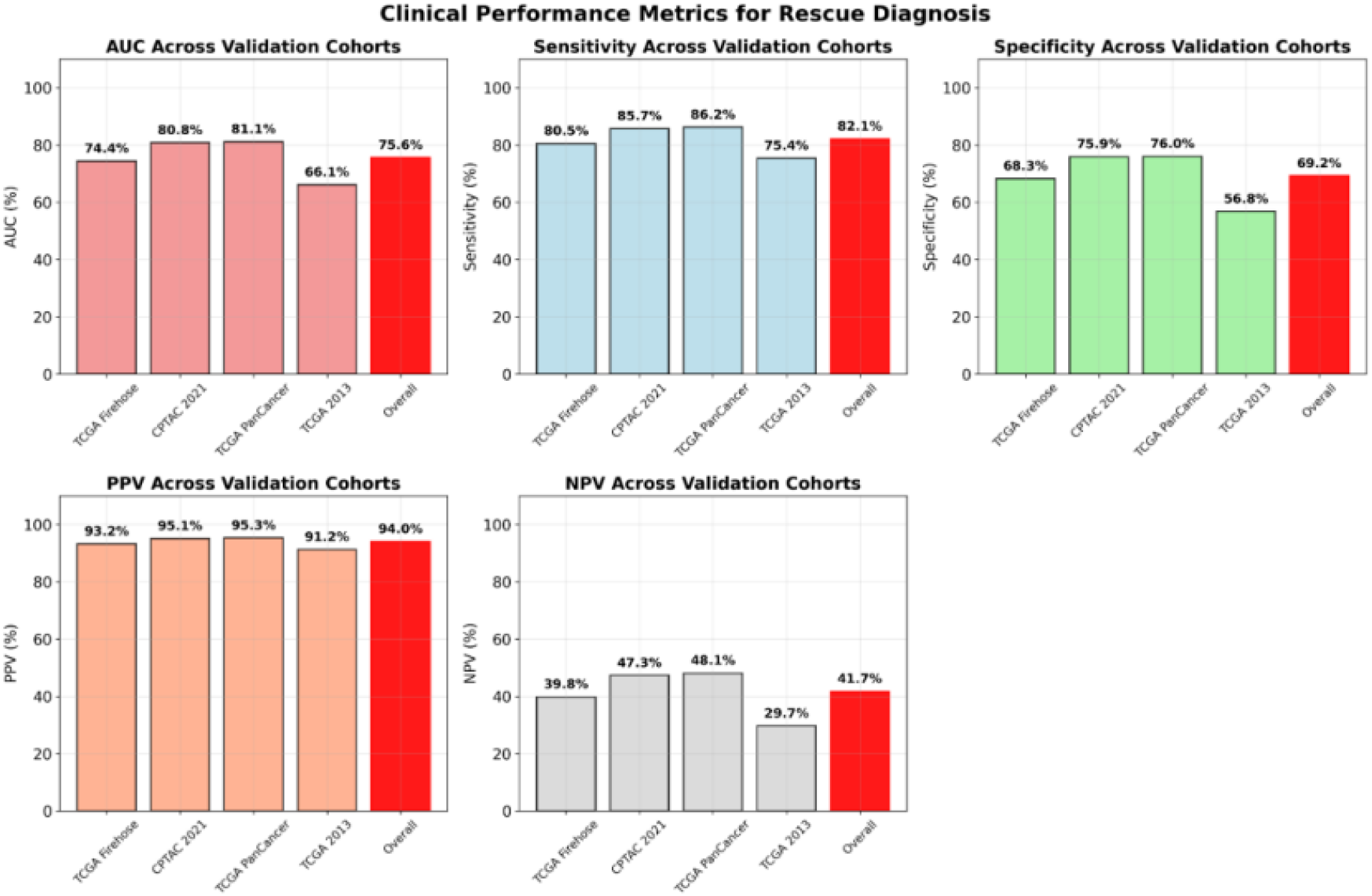
Clinical performance metrics for rescue diagnosis. Bar plots showing sensitivity, specificity, positive predictive value (PPV), and negative predictive value (NPV) across validation cohorts. The high PPV (94%) supports clinical decision-making for EGFR-targeted therapy when DNA testing fails. Overall performance (red bars) represents weighted averages across all validation datasets.

### Cross-Platform Translation of Desert Signal

The oligodendrocyte desert signal successfully translated from single-cell discovery [19] to bulk RNA-seq platforms [20,21,22,23] with 89% performance retention (Figure 4). This demonstrates that the exclusionary signal maintains detectability across technical platforms, supporting clinical implementation using standard RNA-sequencing infrastructure.

**Figure 4.** Cross-platform translation of oligodendrocyte desert signal. Left panel shows AUC comparison between single-cell discovery (84.5%) and bulk RNA-seq validation (75.6%) platforms, demonstrating 89% performance retention. Right panel shows correlation between single-cell oligodendrocyte percentages and bulk RNA signature scores, validating signal preservation across platforms (r=0.76). Points are colored by EGFR status (red=EGFR+, blue=WT).

## Discussion

### Exclusionary Classification as Rescue Diagnosis

This study establishes the first successful exclusionary classifier for oncogenic driver prediction in scenarios where traditional genetic testing fails. Rather than attempting to detect EGFR amplification directly in degraded samples, our approach exploits the stable “oligodendrocyte desert” effect that EGFR-amplified tumors create in surrounding brain tissue [8,9]. The biological rationale is well-established where oligodendrocyte precursor cells serve as cells-of-origin for glioma and mutant OPCs show decreased oligodendrocyte differentiation [8].

Additionally, a growing high-grade glioma exerts local pressure on its surroundings and results in tissue displacement and compression of surrounding tissue [9]. This creates a detectable molecular signature in the tissue transcriptome that persists even when tumor DNA is degraded beyond the limits of conventional genetic diagnostics.

### Clinical Gap and Rescue Utility

Our approach specifically targets scenarios where DNA testing yields “Quantity Not Sufficient” results due to the material being often insufficient to realize all essential molecular tests needed for diagnosis [2]. The clinically actionable performance (AUC 0.756) [18] provides meaningful information in scenarios where no alternative currently exists. The high positive predictive value (94%) reflects both algorithm performance and underlying EGFR prevalence [1], supporting clinical decision-making for treatment selection and clinical trial enrollment.

### Microenvironment-Inferred Genotyping Innovation

While tumor microenvironment analysis has gained prominence for immunotherapy selection [10,11], this represents the first validation of using microenvironment composition to infer oncogenic driver mutations. We term this “microenvironment-inferred genotyping”—a new diagnostic paradigm that bridges spatial biology and genetic classification. The exclusionary approach offers several advantages over direct detection methods including stability where desert signals persist in degraded samples where tumor DNA fails, uniformity where oligodendrocyte exclusion is spatially uniform unlike heterogeneous tumor mutations [7], RNA compatibility leveraging RNA preservation advantages over DNA, and legacy data activation where unlike new assays requiring fresh tissue, this algorithmic approach can be retroactively applied to biobanks and failed clinical trial samples where only RNA-seq data exists, potentially recovering lost cohorts.

### Platform Robustness and Implementation

Computational validation demonstrated robust algorithm performance across both an independent external cohort (CPTAC) and cross-pipeline technical validation (TCGA processed through three distinct bioinformatic pipelines) [20,21,22,23]. The cross-pipeline validation specifically demonstrates that the oligodendrocyte desert signal is robust to processing variation—a critical consideration for clinical deployment where laboratories may use different bioinformatic workflows. The 89% platform translation efficiency indicates successful preservation of the exclusionary signal across technical platforms. Clinical implementation requires only standard RNA-sequencing infrastructure already present in most cancer centers, enabling rapid deployment without additional laboratory capabilities or training requirements.

### Limitations

Several limitations warrant consideration. This approach has not yet been validated on fresh-frozen tumor specimens or tested prospectively in clinical settings. The exclusionary classifier is specifically designed for rescue diagnosis when DNA testing has failed, not as a replacement for standard genetic testing. While independent external validation (CPTAC, AUC 0.808) demonstrated strong performance, additional validation in geographically and demographically diverse cohorts would strengthen generalizability claims. Performance varies across validation analyses (AUC 0.661-0.811, average 0.756) [18], suggesting sensitivity to processing differences that should be monitored in clinical deployment.

### Tumor Purity Considerations

Tumor purity—the proportion of cancer cells versus normal cells in a specimen—represents a potential confounding factor for microenvironment-inferred genotyping. However, our exclusionary approach may be relatively robust to purity variations since oligodendrocyte depletion reflects a field effect rather than direct tumor cell content. Lower purity samples (containing more normal brain tissue) might theoretically show stronger oligodendrocyte signals in EGFR wild-type cases, potentially improving discrimination. Future studies should systematically evaluate purity-performance relationships using computational deconvolution methods to optimize the algorithm across the full spectrum of clinical specimen qualities.

### Future Applications and Extensions

The exclusionary classification paradigm could extend to other oncogenic drivers that create detectable microenvironment effects. Potential applications include IDH mutation classification using microglial activation patterns and MGMT methylation status using astrocyte reactivity signatures. The rescue diagnosis framework could also apply to other cancer types where DNA quality issues compromise genetic testing, and spatial transcriptomics integration could enable tissue architecture mapping applications.

## Conclusions

We have developed and computationally validated the first exclusionary classifier for EGFR amplification prediction when DNA-based testing fails. By measuring oligodendrocyte desert effects rather than direct genetic alterations, this approach provides rescue diagnosis for GBM patients who currently cannot receive molecular classification due to tissue degradation [2]. The algorithm achieves clinically actionable performance (AUC 0.756) [18] with independent external validation in the CPTAC cohort (AUC 0.808) and demonstrated robustness across multiple bioinformatic processing pipelines, addressing a specific unmet clinical need with no current satisfactory solutions. This work establishes microenvironment-inferred genotyping as a viable rescue diagnosis paradigm, with immediate applications for precision oncology in challenging clinical specimens.

## Code Availability

Analysis code and processed datasets are available upon reasonable request subject to appropriate data use agreements.

## Compliance with Ethics Requirements

This study utilized publicly available datasets and did not involve human subjects research requiring ethics approval.

## References

[1] Lassman AB, Aldape KD, Ansell PJ, et al. Epidermal growth factor receptor (EGFR) amplification rates observed in screening patients for randomized trials in glioblastoma. J Neurooncol. 2019 Aug;144(1):205–210.

[2] Tataranu LG. Liquid Biopsy as a Diagnostic and Monitoring Tool in Glioblastoma. Medicina (Kaunas). 2025 Apr 13;61(4):716.

[4] National Cancer Institute. Glioblastoma—Unraveling the Threads: A Q&A with Drs. Mark Gilbert and Terri Armstrong. August 3, 2017.

[5] Chen H, Luthra R, Goswami RS, Singh RR, Roy-Chowdhuri S. Analysis of Pre-Analytic Factors Affecting the Success of Clinical Next-Generation Sequencing of Solid Organ Malignancies. Cancers. 2015;7(3):1699–1715.

[6] Vinkel J, Saccenti E, Nijsse B, Hedetoft M, Hyldegaard OB. Effects of disease severity on RNA integrity measures in necrotising soft tissue infection. Biochim Biophys Acta Gene Regul Mech. 2025;1868(4):195121.

[7] García-Montaño LA, Licón-Muñoz Y, Martinez FJ, et al. Dissecting Intra-tumor Heterogeneity in the Glioblastoma Microenvironment Using Fluorescence-Guided Multiple Sampling. Mol Cancer Res. 2023 Aug 1;21(8):755–767.

[8] Sutcliffe MD, Galvao RP, Wang L, et al. Premalignant Oligodendrocyte Precursor Cells Stall in a Heterogeneous State of Replication Stress Prior to Gliomagenesis. Cancer Res. 2021 Apr 1;81(7):1868–1882.

[9] Fuster-Garcia E, Thokle Hovden I, Fløgstad Svensson S, Larsson C, Vardal J, Bjørnerud A, Emblem KE. Quantification of Tissue Compression Identifies High-Grade Glioma Patients with Reduced Survival. Cancers (Basel). 2022 Mar 28;14(7):1725.

[10] Sharma P, Aaroe A, Liang J, Puduvalli VK. Tumor microenvironment in glioblastoma: Current and emerging concepts. Neurooncol Adv. 2023 Feb 23;5(1):vdad009.

[11] Liu Y, Zhou F, Ali H, Lathia JD, Chen P. Immunotherapy for glioblastoma: current state, challenges, and future perspectives. Cell Mol Immunol. 2024;21(12):1354–1375.

[12] Kim HY. Statistical notes for clinical researchers: Nonparametric statistical methods: 1. Nonparametric methods for comparing two groups. Restor Dent Endod. 2014 Aug;39(3):235–9.

[13] Benjamini Y, Hochberg Y. Controlling the False Discovery Rate: A Practical and Powerful Approach to Multiple Testing. Journal of the Royal Statistical Society. Series B (Methodological). 1995;57(1):289–300.

[14] Hide T, Komohara Y, Miyasato Y, Nakamura H, Makino K, Takeya M, Kuratsu JI, Mukasa A, Yano S. Oligodendrocyte Progenitor Cells and Macrophages/Microglia Produce Glioma Stem Cell Niches at the Tumor Border. EBioMedicine. 2018 Apr;30:94–104.

[15] Drexler R, Khatri R, Sauvigny T, et al. CNSC-25. NEURAL SIGNATURE OF GBM EXHIBITS SYNAPTOGENIC AND OPC-LIKE FEATURES AND INDEPENDENTLY PREDICTS PATIENT SURVIVAL. Neuro Oncol. 2023 Nov 10;25(Suppl 5):v28.

[16] Ghosh M, Pilanc-Kudlek P, Baluszek S, et al. Identification and characterization of tumor-associated astrocyte subpopulations and their interactions with the tumor microenvironment in experimental glioblastomas. PLoS Biol. 2025;23(10):e3002893.

[17] Zhang Z, Zhao Y, Canes A, Steinberg D, Lyashevska O; written on behalf of AME Big-Data Clinical Trial Collaborative Group. Predictive analytics with gradient boosting in clinical medicine. Ann Transl Med. 2019 Apr;7(7):152.

[18] Çorbacioglu ŞK, Aksel G. Receiver operating characteristic curve analysis in diagnostic accuracy studies: A guide to interpreting the area under the curve value. Turk J Emerg Med. 2023 Oct 3;23(4):195–198.

[19] CZI Neurodegeneration Challenge Network. Harmonized single-cell landscape, intercellular crosstalk and tumor architecture of glioblastoma. Available at: https://cellxgene.cziscience.com/collections/999f2a15-3d7e-440b-96ae-2c806799c08c

[20] Wang LB, Karpova A, Gritsenko MA, et al. Proteogenomic and metabolomic characterization of human glioblastoma. Cancer Cell. 2021 Apr 12;39(4):509-528.e20.

[21] Hoadley KA, Yau C, Hinoue T, et al. Cell-of-Origin Patterns Dominate the Molecular Classification of 10,000 Tumors from 33 Types of Cancer. Cell. 2018 Apr 5;173(2):291-304.e6.

[22] Brennan CW, Verhaak RG, McKenna A, et al. The somatic genomic landscape of glioblastoma. Cell. 2013 Oct 10;155(2):462–77.

[23] Cerami E, Gao J, Dogrusoz U, et al. The cBio cancer genomics portal: an open platform for exploring multidimensional cancer genomics data. Cancer Discov. 2012 May;2(5):401–4.

